# Genome-wide diversity and differentiation of two novel multidrug-resistant populations of *Pasteurella multocida* type B:2 from fowl cholera

**DOI:** 10.1101/2020.08.24.262618

**Authors:** Otun Saha, M. Rafiul Islam, M. Shaminur Rahman, M. Nazmul Hoque, M. Anwar Hossain, Munawar Sultana

## Abstract

*Pasteurella multocida* is the etiologic agent of fowl cholera (FC), a highly contagious and severe disease in poultry with higher mortality and morbidity. Twenty-two *P. multocida* strains isolated from the FC outbreaks were subjected to phenotypic and genotypic characterization. The isolates were grouped into two distinct RAPD biotypes harboring a range of pathogenic genes; *exb*B, *omp*H, *ptf*A, *nan*B, *sod*C, and *hgb*A. Among these strains, 90.90% and 36.37% were multidrug-resistant and strong biofilm formers, respectively. Whole genome sequencing of the two representative RAPD isolates confirmed as *P. multocida type* B:L2:ST122 harboring a number of virulence factors, and antimicrobial resistance genes. Pan-genome analysis revealed 90 unique genes in these genomes associated with versatile metabolic functions, pathogenicity, virulence, and antimicrobial resistance. This study for the first time reports the association of *P. multocida* genotype B:L2:ST122 in the pathogenesis of FC, and provides a genetic context for future researches on *P. multocida* strains.

## 1. Introduction

Poultry raring is one of the important sources of income in Bangladesh, including both commercial broiler, and layer farming. This subsector of livestock has been noted as the largest primary source of eggs, and meat since last decade, and contributes ~3.0% to the GDP of Bangladesh [1]. However, this industry faces a number of constraints including limited feed resource, and frequent outbreaks of infectious diseases. Emerging infectious diseases with zoonotic potentials are appearing at an alarming rate, and pose a growing threat to global biodiversity [2, 3]. With this backdrop, there are rising apprehensions among health care providers, veterinarians, policymakers, and the general public about human acquisition of zoonotic infections from nearby encounters with domesticated pets, wild animals, and/or avian species like commercial poultry [4–6].

*Pasteurella multocida* is identified as a major bacterial etiologic agent for many infectious diseases in a wide spectrum of domestic and wild animals such as poultry and wild birds, pigs, cattle and buffaloes, rabbits, small ruminants, cats, dogs and other mammals [4, 7–9]. *P. multocida* is a zoonotic Gram negative, and opportunistic bacterium [10], and causes FC or pasteurellosis, an economically important disease to the poultry industry [8, 11]. Among the bacterial diseases, FC is a major threat to the poultry industry causing 25-35% mortality [12]. Different strains of *P. multocida* are associated with clinical manifestations of FC ranging from depression, anorexia, lameness, swollen wattles, mucoid discharge from the mouth, ruffled feathers, diarrhea, increased respiratory rate, and per acute or sudden asymptomatic death [12, 13]. The disease can range from acute septicemia to chronic and localized infections, and the morbidity and mortality may be up to 100%. The route of infection is oral or nasal with transmission via nasal exudate, faces, contaminated soil, equipment, and people [14].

Currently, the strains of *P*. *multocida* are divided into 5 serotypes such as A (*hya*D*-hya*C), B (*bcb*D), D (*dcb*F), E (*ecb*J), and F (*fcb*D based on capsular typing [7, 8]. Among these serotypes, capsular serogroups A and F cause the majority of FC, whereas serogroups B and E are predominantly associated with hemorrhagic septicemia (HS) in cattle and wild ruminants [4, 15]. In addition to capsular serogroups, *P. multocida* strains are currently classified into 16 Heddleston lipopolysaccharide (LPS) serovars [16, 17]. Different serotypes of the *P. multocida* exhibit varying degrees of virulence in different host [7, 8]. Various strains of *P. multocida* harbor a wide arsenal virulence factors-associated genes (VFGs), and of them, genes involved in the capsule formation, LPS, fimbriae and adhesins, toxins, iron regulation and acquisition proteins, sialic acid metabolism, hyaluronidase, and outer membrane proteins (OMPs) are the key components of regulating the pathogenesis [4, 7, 18, 19]. Though not well studied, many of these VFGs might play a substantial role in the pathogenesis of FC, and survival in the complex host environment [7, 18, 19]. Previous reports showed that there is a clear correlation between certain VFGs, and capsular types or biovars [20, 21]. Therefore, identification of prevalent VFGs is important to predict the pathogenic nature of the bacterium, and select potential vaccine candidates.

Till to date, there is lack of effective coverage in multi-serotype vaccines, and thus, antibiotics remain as the most commonly employed treatment strategies for prevention and control of FC despite their increased correlation to drug resistance, and food safety risks due to residual effects [4, 8, 22]. Furthermore, the infrequent, and unnecessary overuse of antibiotics in the poultry farms, livestock, fisheries, and agriculture is believed to have augmented the emergence of MDR pathogenic bacterial species like *P. multocida* posing a serious threat to public health and livestock [4].

Bio-molecular techniques such as polymerase chain reaction (PCR), ribotyping, random amplification of polymorphic DNA (RAPD)-PCR, multilocus sequence type (MLST), and 16S rRNA gene sequencing analysis enabled the studies of molecular epidemiology, and genetic diversity of *P. multocida* [18, 23–25]. In addition to these molecular techniques, whole-genome sequencing (WGS) is an affordable, convenient, and rapid technique for outbreak investigations, diagnostics, and epidemiological surveillance [26, 27]. The state of the art WGS technology might enable us to study the underlying genetic mechanisms associated with pathogenicity, virulence fitness, and host adaptability of *P. multocida* in multiple hosts [4, 28, 29]. However, there is limited information on the molecular epidemiology of *P. multocida* circulating in the global poultry industry with particular reference to Bangladesh. Therefore, in this study, we’ve employed different bio-molecular techniques along with the WGS of two representative MDR, and SBF *P. multocida* isolates to investigate the phenotypic and genotypic diversity, and underlying genetic contents such as VFGs, antimicrobial resistance genes (ARGs), and metabolic functional potentials to be associated with the pathogenesis of FC in laying hens in Bangladesh. Moreover, herein this study, we first time report the phenotypic and genotypic traits, and associated genetic factors of highly pathogenic strains of *P. multocida* type B:2 causing FC in chickens of Bangladesh.

## 2. Methods

### 2.1. Sample collection, bacterial strain isolation, and genomic DNA extraction

We collected fifty-seven samples (n=57) of suspected FC including 36 suspected live birds, and 21 dead laying hens from six commercial layer farms located in Narsingdi (23.9193° N, 90.7176° E), Narayangonj (3.6238° N, 90.5000° E), and Manikgonj (23.8617° N, 90.0003° E) districts of Bangladesh during August 2017 to November 2017. Diseased birds were diagnosed with FC by observing clinical signs and symptoms including sudden death, swollen wattle and combs, lameness, respiratory rales, and diarrhea by practicing veterinarians. The birds were then dissected, and internal organs (liver) from each bird were collected as the experimental samples. The samples were finally processed, kept in the nutrient enriched media, and transported to the laboratory (at 4°C). The samples were plated on Luria-Bertani broth (LB) (Oxoid, Thermo Fisher Scientific, UK). A small amount of inoculum from LB was streaked onto Blood agar based (BAB) (Oxoid, Thermo Fisher Scientific, UK) supplemented with 5% sheep blood, and incubated for 24 h at 37 °C for selective growth. Colonies revealing the characteristic of the *P. multocida* on the BAB agar (3–5 colonies from each sample) were further inoculated into MacConkey (MC) agar. The MC agar positive colonies were further subjected to biochemical tests according to Kim et al. (2019) [7]. A total of 78 bacterial isolates were retrieved from the selective culture and biochemical tests, and of them 22 isolates were confirmed as *P. multocida* by *kmt1* gene-specific PCR following previously published protocols [30]. Genomic DNA of *P. multocida* was extracted from overnight culture by the boiled DNA extraction method [31]. The quality and quantity of the extracted DNA was measured using a NanoDrop ND-2000 spectrophotometer (Thermo Scientific, Waltham, USA). The extracted DNA was kept at −80 °C [32].

### 2.2. Molecular typing and detection of pathogenic genes

Random amplification of polymorphic DNA (RAPD)-PCR was performed with the extracted DNA to investigate the biotype diversity among the isolates [33, 34]. The RAPD-PCR was performed by using (5’-GCGATCCCCA-3’) primer following the previously optimized protocol for *E. coli* [31]. Using previously published pathogenic gene-specific primers (Table 1) for PCR assays, we surveyed 10 pathogenic genes in the isolated strains of *P. multocida* [7, 19]. Briefly, each PCR reaction contained 2 μL DNA template (300 ng/μL), 10 μL PCR master mix 2X (Go Taq Colorless Master Mix) and 1μL (100 pmol/μL) of each primer (Table 1) in each tube. The PCR amplifications were conducted in thermocycler, and the cycling conditions were identical for all the samples as follows: 94 °C for 5 min; 35 cycles of 1min at 94 °C, 1 min at 50-60 °C, and 1 min at 72 °C; and 72 °C for 7 min. PCR amplicons were visualized on 1.5% agarose gel prepared in 1× TAE buffer. After gel electrophoresis, the images were captured using Image ChemiDoc™ Imaging System (Bio-Rad, USA) [7].

**Table 1.**
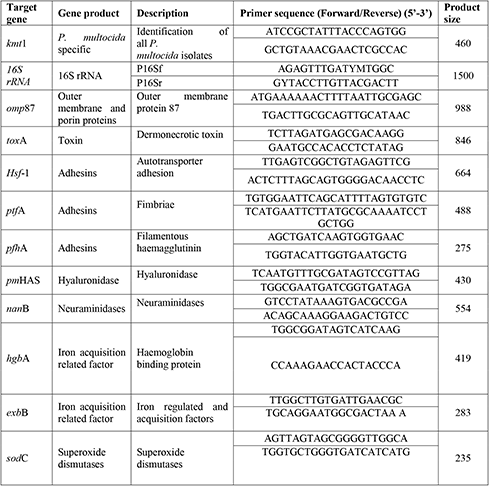
Primers for genotyping and pathogenic gene (VFGs) characterization in *P. multocida* strains.

### 2.3. Antimicrobial susceptibility testing

The antimicrobial susceptibility testing of various *P. multocida* pathotypes was carried out using Kirby-Bauer disc diffusion method on Mueller-Hinton agar according to the guidelines of the Clinical and Laboratory Standards Institute; CLSI [35], and European Committee on Antimicrobial Susceptibility Testing; EUCAST [36]. Antibiotics were selected for susceptibility testing corresponding to a panel of antimicrobial agents of interest to the poultry industry and public health in Bangladesh. The *kmt*1 gene positive pathogenic isolates (n=22) were selected for antimicrobial susceptibility to 17 antibiotic discs comprising ampicillin (AMP) (10 mg), oxacillin (OXA) (10mg), doxycycline (DOX) (30 mg), streptomycin (STR) (10 mg), nitrofurantoin (F) (300 mg), levofloxacin (LEV) (3^rd^generation) (5 mg), aztreonam (ATM) (30 mg), cefoxitin (Fox) (30 mg), gentamycin (CN) (10 mg), nalidixic acid (NA) (1st generation) (30 mg), trimethoprim (Tm) (5 mg), tetracycline (TE) (30 mg), ciprofloxacin (CIP) (2^nd^ generation) (5 mg), chloramphenicol (C) (30 mg), sulfonamide (S3) (250μg), colistin sulfate (CS) (10mg), and cefepime (FEP) (30 mg). Finally, the findings were recorded as susceptible, intermediate and resistant according to the CLSI and EUCAST breakpoints.

### 2.4. Biofilms formation (BF) assay

The BF ability of the *P. multocida* pathotypes was tested in 24-well polystyrene plates (Corning, Costar) as previously described [37]. Briefly, all isolated (n=22) *P. multocida* strains were grown in LB medium for 24□h at 37□°C with shaking, and a 1:1000 dilution was prepared in LB. Twenty-five microliters were placed in each well containing 1.5□ml of culture medium. The plates were incubated for 48□h at 37□°C in static condition. Planktonic cells were removed, and wells containing biofilms were rinsed three times with distilled water, and the remaining adherent bacteria were stained with 2□ml/well of crystal violet (0.7% [wt/vol] solution; Sigma-Aldrich) for 12□min. Excess stain was removed by washing with distilled water. Crystal violet (CV) was extracted by acetic acid (33% [vol/vol]), and the plates were incubated at room temperature to release the dye into the solution. Then, two samples of 100□μl from each well were transferred to a 96-well flat-bottom plate, and the amount of dye was determined at 600□nm using a microplate reader. As quantified in crystal violet assay, the bacterial strains were divided into four groups based upon optical density (OD) at 600 nm of the bacterial biofilm: non-biofilm former (NBF), weak biofilm former (WBF), moderate biofilm former (MBF) and strong biofilm former (SBF) [37].

### 2.5. Ribosomal gene (16S rRNA) sequencing and phylogenetic analysis

Two *P. multocida* isolates; PM4 and PM7, representative of each genotype and/or biotype, were selected for ribosomal gene (16S rRNA) sequencing using universal primers (Table 1) followed by sequencing at First Base Laboratories SdnBhd (Malaysia) using Applied Biosystems highest capacity-based genetic analyzer (ABI PRISM^®^ 377 DNA Sequencer) platforms with the BigDye^®^ Terminator v3.1 cycle sequencing kit chemistry [38]. Initial quality controls of the generated raw sequences were performed using SeqMan software, and were aligned with relevant reference sequences retrieved from NCBI database using Molecular Evolutionary Genetics Analysis (MEGA) version 7.0 for bigger datasets [39]. We employed the Kimura-Nei method to compute the evolutionary distances, and neighbor-joining method to construct a phylogenetic tree [40]. The percentage of replicate trees in which the associated taxa clustered together in the bootstrap test (1000 replicates) is shown next to the branches. The evolutionary distances were computed using the Kimura 2-parameter method and in the units of the number of base substitutions per site [41].

### 2.6. Whole genome sequencing and assembly

Two RAPD representative *P. multocida* pathotypes (PM4 and PM7) were selected for whole-genome sequencing (WGS). Genomic DNA libraries containing 300∼500 bp fragments were constructed using Nextera XT DNA Library Preparation Kit (Illumina Inc., San Diego, USA). The WGS was performed using 100 bp paired-end sequencing protocol under Illumina platform using HiSeq4000 sequencer (Macrogen, lnc. Seoul, Republic of Korea) with an average ∼456-fold genome coverage per sample. The generated FASTQ files were evaluated for quality using FastQC v0.11 [42]. Adapter sequences, and low-quality ends per read were trimmed by using Trimmomatic v0.39 [43] with set criteria of sliding window size 4; a minimum average quality score of 20; minimum read length of 50 bp. High quality reads were *de novo* assembled to generate contigs using an open source platform, SPAdes (Species Prediction and Diversity Estimation) v3.13 [44] with a range of k-mer between 21 to 121, and contigs less than 500 bp were filtered. The draft contigs were mapped, reordered, and scaffolded according to a NCBI reference complete sequence of *P. multocida* strain Razi_Pm0001 (accession number: NZ_CP017961.1) by RaGOO v1.1 [45]. Completeness of the scaffold was checked using CheckM [46]. Scaffolded contigs were searched for the bacterium at strain level by BLAST, and the k-mer algorithm in the KmerFinder 3.1 tool [47]. The WGS-based phylogenetic tree was constructed using the online pipeline Reference Sequence Alignment Based Phylogeny Builder (REALPHY) [48], and visualized on iTor v5 [49]. The Plasmid Finder v2.1, and Integron_Finder v1.5 tools were used for the detection of plasmid sequence contamination [50, 51]. Prophage sequences within the genome assemblies were identified by the PHAge Search Tool Enhanced Release (PHASTER) web server [52].

### 2.7. Genome annotation and genomic organization mapping

The scaffolded genomes of the *P. multocida* PM4 and PM7 strains were annotated by multiple annotation schemes to improve accuracy. We used the NCBI Prokaryotic Genome Annotation Pipeline (PGAPv4.11) with best-placed reference protein set, and GeneMarkS-2+ annotation methods [53], Rapid Prokaryotic Genome Annotation (Prokka) (e = 0.000001) [54], and Rapid Annotation using Subsystem Technology (RAST) (e = 0.000001) [55]. Annotated genes by each tool were then cross-checked. To remove the tRNA and mRNA from the genomes, we used tRNAscan-SE v2.0 [56], and Aragorn v1.2.38 [57] tools. The graphical map of the circular genome was generated using the CGView Server (http://stothard.afns.ualberta.ca/cgview_server/) [58]. Circular BLAST of annotated genome comparison was performed using an in-house instance of BLAST Ring Image Generator (BRIG) v0.95 [59]. The pan-genome and core-genome were analyzed using the Bacterial Pan Genome Analysis (BPGA) pipeline [60].

### 2.8. Genome sequence analysis

The trimmed fastq files and assembled genomes of PM4 and PM7 were aligned with *P. multocida* capsular serotypes (*hyaD-hyaC, bcbD, dcbF, ecbJ, fcbD*) [7]; LPS genotypes (LPS outer core structure genes) [17] genes using Burrows-Wheeler Aligner (BWA) [61], and the output was extracted using Resistome Analyzer integrated into AmrPlusPlus v2.0 pipeline [62]. MLST genotypes of the *P. multocida* strains were assigned by performing blast of their scaffolds against the *P. multocida* MLST Databases (https://pubmlst.org/pmultocida). In this study, RIRDC MLST, a scheme of multi-locus sequence typing based on seven housekeeping genes (*adk, est, pmi, zwf, mdh, gdh*, and *pgi*) of *P. multocida* in the Bacterial Isolate Genome Sequence Database (BIGSdb) [63] was used to determine the MLST genotypes of the isolates (PM4 and PM7).

### 2.9. Virulence factors (VFGs), antimicrobial resistance (ARGs) and metabolic functions profiling

The trimmed fastq files, and assembled genomes of PM4 and PM7 were further aligned with *P. multocida* outer membrane and porin proteins (*oma*87*, omp*H*, plp*B*, psl*), adesins (*ptf*A*, fim*A*, hsf-1, hsf-2, pfh*A*, tad*D); neuraminidases (*nan*B*, nan*H), iron acquisition related factors (*exb*D*, ton*B*, fur, tbp*A*, hgb*A*, hgb*B); superoxide dismutases (*sod*A*, sod*C); dermonecrotoxin *(tox*A); and hyaluronidase (*pm*HAS) pathogenic genes using BWA [61], and the resulting output was using Resistome Analyzer integrated into AmrPlusPlus v2.0 pipeline [62]. We mapped the annotated genomes against the ResFinder Database [64], Comprehensive Antibiotic Resistance Database (CARD) [65], MEGARes [62, 66] databases to search for ARGs, and the gene fraction value ≥ 95 was considered ARGs to our strains. Assembled genomes were also blasted against Virulence Factor database (VFDB) [67] for putative VFGs identification. We predicted the metabolic functional potentials of the two genomes through the KEGG (Kyoto Encyclopedia of Genes and Genomes) Automatic Annotation Server (KAAS) [68] database.

## 3. Results and Discussion

### 3.1. Intra-outbreak diversity of P. multocida in FC

The present study focused on the phenotypic and genotypic characterization of pathogenic *P. multocida* associated with fowl cholera in commercial layer birds. We obtained 78 *P*. *multocida* isolates from 57 samples originating from different layer farms of the Narsingdi, Narayangonj and Manikgonj districts of Bangladesh. The isolates produced small glistening dewdrop like colonies on blood agar plates after incubation at 37 °C for 18 h, and appeared as Gram-negative cocco-bacilli with Gram’s staining. Other characteristics cultural properties included no growth on MacConkey agar, and bacteria remained non-motile and non-hemolytic on blood agar. Of the retrieved isolates (n = 78), the species and type-specific PCR using *kmt*1 gene confirmed 22 isolates as *P. multocida* from the samples of Narsingdi district (Table 2, Fig. S1), which was further confirmed through biochemical and molecular characterization [7, 30]. Since the isolates from two other district, Narayangonj and Manikgonj lacked *kmt*1 gene, these might include different bacterial species, which require further investigations. The *kmt*1 gene positive 22 isolates of *P. multocida* were catalase and oxidase positive, and urease negative. Furthermore, there was no reaction with citrate, methyl red, and Voges-Proskauer (Table 2). These isolates fermented glucose, mannose and sucrose, and none utilized lactose. However, fermentation of inositol grouped these *P. multocida* isolates into two biotypes (Table 2). Out of 22 *P. multocida* isolates, 14 (63.64%) isolates fermented inositol, and denoted as biotype 1 (Table 2). Conversely, 8 (36.36%) isolates remained unable to ferment inositol, and thus denoted as biotype 2 (Table 2). These results indicated that *P. multocida* isolates might possess diverse metabolic potentials despite being identified from the same outbreak of FC as also reported earlier [22]. Moreover, the RAPD assay also confirmed two pathotypes of *P. multocida* among these 22 strains (Table 2, Fig. S2). In this study, the RAPD profiling indicated less genetic heterogeneity among the studied *P. multocida* strains corroborating the findings of Hotchkiss et al. [69]. There we found a positive correlation between biotypes and RAPD profiles since all of the 14 isolates of biotype 1 showed the RAPD pattern 1 while the rest of the 8 isolates belonged to biotype 2 showed RAPD pattern 2 (Table 2). Moreover, pathogenic gene-specific primer-based PCR revealed different pathogenic genes among the identified strains (Table 2). Three pathogenic genes *nan*B, *sod*C, and *hgb*A were found in all of the tested isolates, and 22.73% (5/22) isolates harbored six pathogenic genes (*exb*B, *omp*H, *ptf*A, *nan*B, *sod*C, and *hgb*A) out of the 10 VFGs tested (Table 2). The present results therefore show that these VFGs are highly prevalent in FC associated *P. multocida* isolates indicating their high pathogenic potentials. Although the molecular basis of the pathogenicity and host specificity of *P. multocida* is not well understood, several studies have reported that a number of VFGs are correlated with the pathogenic mechanisms of FC [7, 14, 22]. However, corroborating the previous observations by Omaleki et al. [29], the biochemical and molecular findings of the current study reflect the phenotypic and genotypic diversity of the *P. multocida* isolates originated from the same outbreak of FC.

**Table 2.**
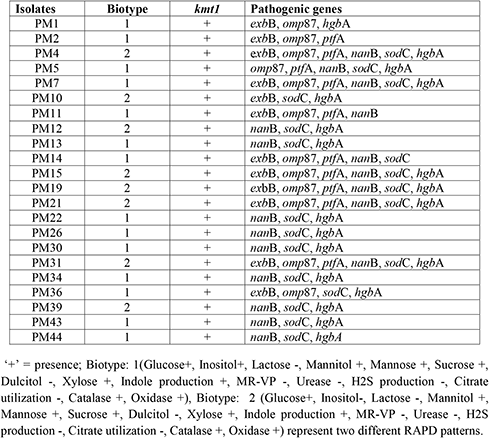
The bio-molecular features, and VFGs profile of the *P. multocida* isolates (n=22).

### 3.2. Antimicrobial resistance in FC-associated P. multocida strains

Antibiotic susceptibility of the recovered *P. multocida* strains (n=22) was tested against 17 different antibiotics of varied classes through disc diffusion assay. The prevalent strains detected were resistant to ampicillin (90.91%), tetracycline (90.91%), and nalidixic acid (63.64%) according to the EUCAST breakpoints (Fig. 1). In this study, 68.18% *P. multocida* strains found to be resistant against each of the three antibiotics, oxacillin, cefoxitin and trimethoprim. Remarkably, we observed that 91.91% isolates were multidrug-resistant, being resistant against ≥5 antibiotics tested (Fig. 1). Treatment for infections with *P. multocida* either in livestock or in poultry with FC or pasteurellosis commonly includes broad-spectrum antimicrobials. The findings of our antimicrobial resistance (AMR) study, like the findings of Jabeen et al. [25] in Malaysia, Kehrenberg et al. [70] in France, Yoshimura et al. [71] in Japan, and Tang et al. [72] in China indicated that ampicillin, tetracycline, and nalidixic acid were the most resistant antibiotics. Therefore, preventive and therapeutic effects on birds suffering from FC with *P. multocida* strains should no longer be expected from these antibiotics. However, all of the tested isolates were sensitive to colistin, suggesting that this antibiotic could effectively be used for treatment of FC or other infections with *P. multocida*. The results documented that *P. multocida* has developed resistance to commonly used antibiotics in poultry. Frequent and excessive use of antibiotics in the livestock of Bangladesh might have a role in AMR development against multiple antibiotics in the clinical infections caused by *P. multocida* [73, 74].

**Fig. 1.**
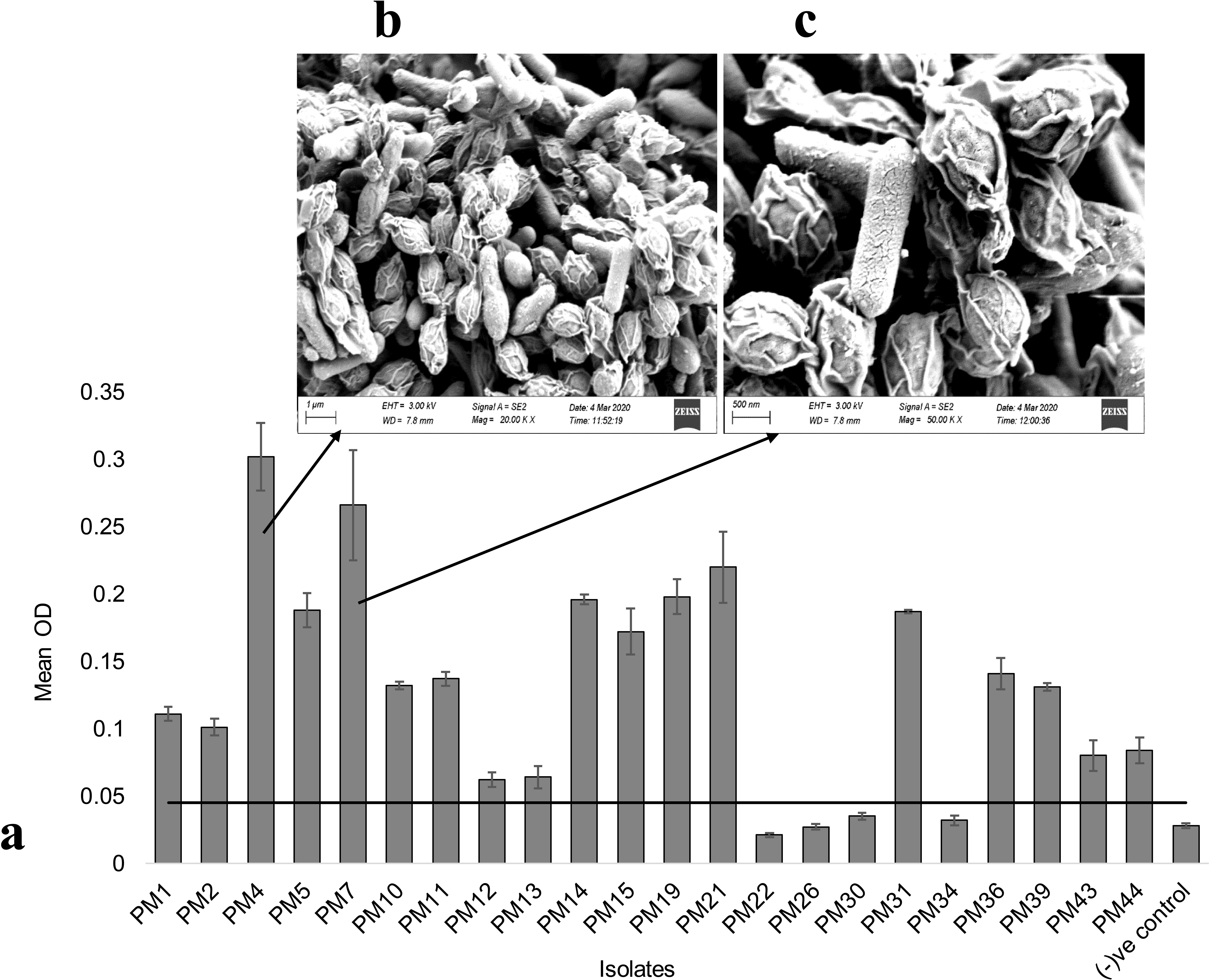
Antimicrobial resistance profile of the tested 22 *P. multocida* strains. Antibiotic susceptibility to 17 antibiotics of varied classes was determined by disk diffusion assays. The strains were categorized as resistant or susceptible based on the breakpoints defined by the European Committee on Antimicrobial Susceptibility Testing (EUCAST, 2020). Numeric inside the gray boxes represent zone of inhibition (in mm unit) for the antibiotics that have no standard breakpoints currently available. Biofilm producing abilities are shown in the right column based on their adherence potential on 24-well polystyrene plates. The superscript asterisks (*) in two isolates (PM4 and PM7) indicate that they were selected for WGS.

### 3.3. Biofilm formation infers higher antibiotic resistance (ABR) in P. multocida

Among the mechanisms of bacterial virulence, biofilm is recognized as an intricate factor in many bacterial infections, and one of the reasons for treatment failure with antibiotics [74–76]. However, in contrast to other microorganisms, only few studies have examined the of biofilm formation (BF) properties of the *P. multocida* [75]. Moreover, studies comparing the pathogenicity of *P. multocida* strains as a function of their biofilm production capacity are also rare in avian species. The *P. multocida* strains isolated from FC formed biofilms on polystyrene plates, as shown in Fig. 2. In this study, the average OD of the negative control was 0.028±0.002, and the cutoff OD value was set as 0.045. The isolates which have OD value ≤0.045 were considered as NBF. In this study, 81.82% (18/22) of the *P. multocida* strains were biofilm-formers with significance differences (P = 0.039) in their BF categories, and we denoted them as strong biofilm former (SBF, 36.36%), moderate biofilm former (MBF, 27.27%), and weak biofilm former (WBF, 18.18%). In our present study, 18.18% of study strains did not produce biofilms, and thus, designated as the non-biofilm formers (NBF) (Fig. 2a) In addition, the SBF isolates showed higher AMR properties (41.18-64.71%) compared to the MBF (14.50-34.65%), and WBF (5.88-29.91%) strains against the tested antibiotics. Moreover, 35.29 - 41.18% of the NBF strains also showed AMR phenomena against these antibiotics (Fig. 1). Furthermore, scanning acoustic microscopy (SAM) of the two representative *P. multocida* strains (Isolate: PM4 and PM7), as representative of SBF strains demonstrated densely colonization, and exopolysaccharides covering the bacterial cells, validating the high biofilm potential of these two MDR strains (Fig. 2b, c). Results from antibiogram, and BF assays indicate that BF ability of the 0.*P. multocida* may enhance antibiotic resistance and pathogenic fitness of *P. multocida* to survive under unfavorable complex conditions within host and environmental niches [76]. Moreover, the biofilms may also promote the bacterium to resist host immune defense mechanisms [74]. Therefore, BF of *P. multocida* needs to be studied in-depth, which might be an alternative to combat the threat of drug-resistant pathogenic *P. multocida* [77].

**Fig. 2.**
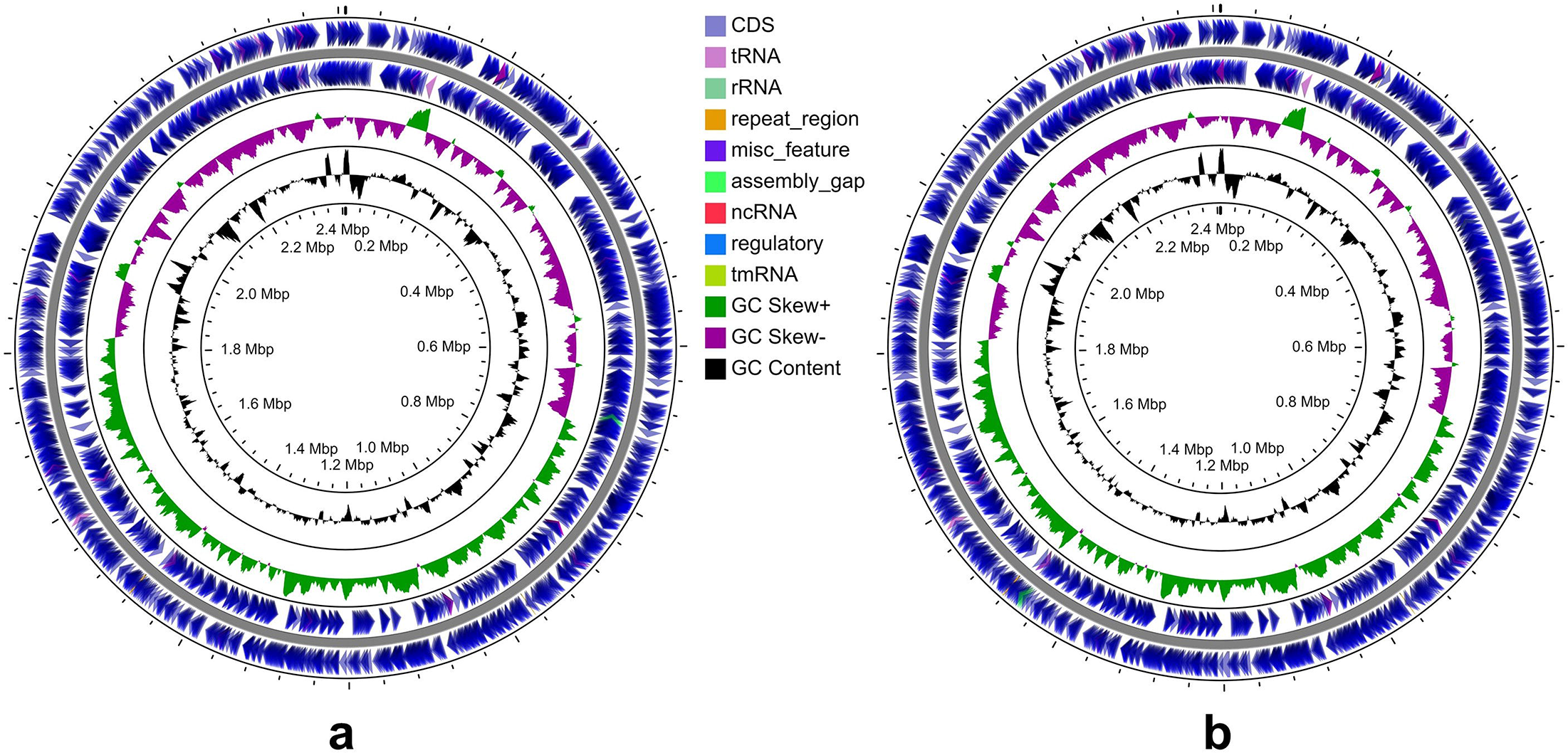
Biofilm formation (BF) ability of the *P. multocida* isolates. (a) Mean optical density (OD) of the 22 isolates measured at 600 nm after 48 h growth at 37 °C. The horizontal line represents the threshold below which indicates non-biofilm producers. Biofilms of two strong biofilm-producing isolates, (b) PM4 and (c) PM7 were further observed under scanning acoustic microscopy (SAM). The SAM micrographs demonstrated the colonizing bacterial cells after 48 h incubation, and revealed how the bacteria tend to grow in clumps (micro-colonies), and the exopolysaccharide that is covering the bacteria. Error bars represent standard deviation.

**Fig. 3.**
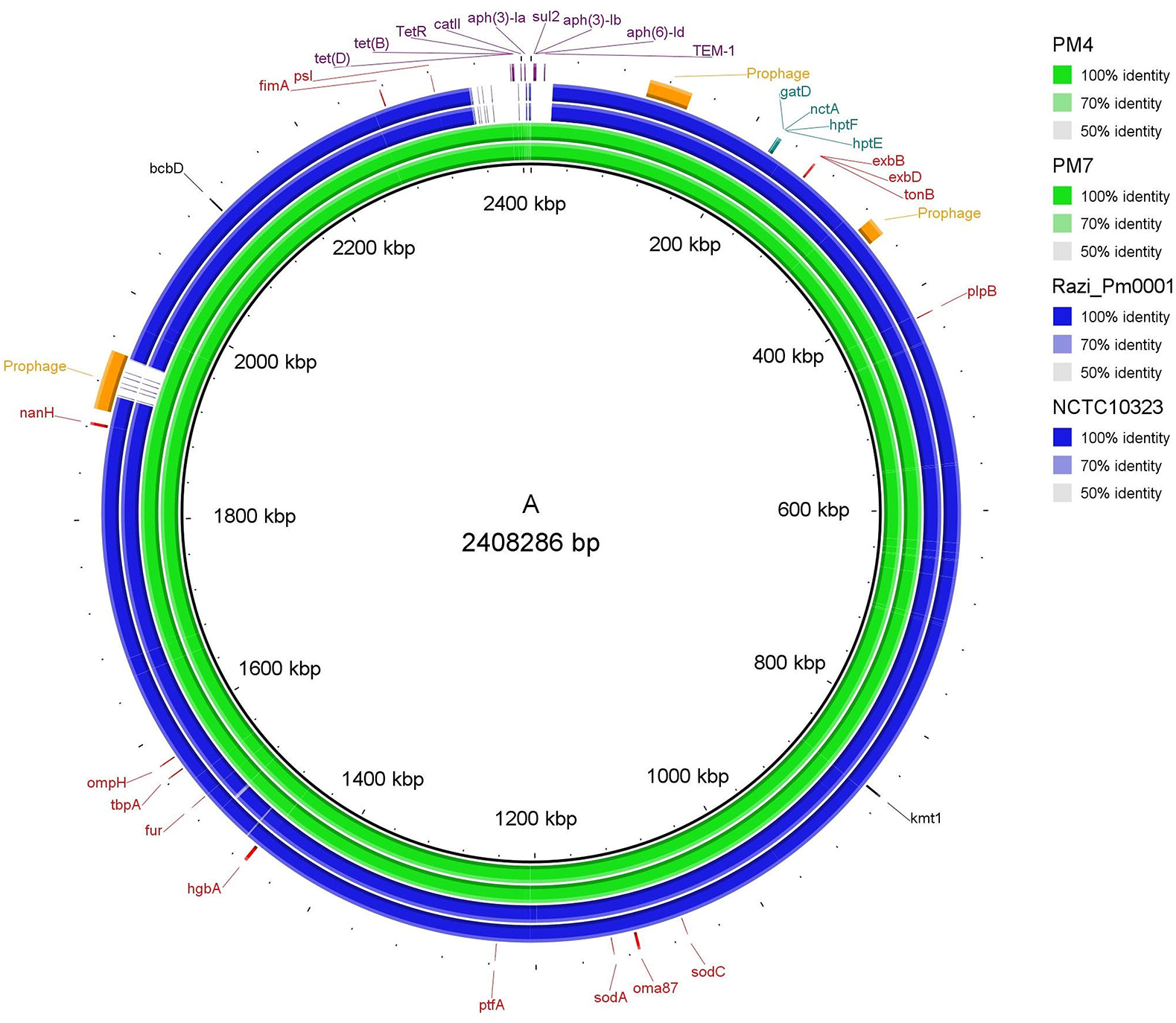
Circular representation of the genome of of *P. multocida* (a) PM4 and (b) PM7 strains. The PM4 and PM7 genomes, and their coding regions with homologies, the tRNA and rRNA operons, and the overall G-C content are presented. The outer two circles demonstrate the coding sequence (CDS), tRNA, and rRNA. The third circle shows the GC content (black). The fourth circle represents the GC skew curve (positive GC skew, green; negative GC skew, violet). The figures were generated by using CGView Server (http://stothard.afns.ualberta.ca/cgview_server/).

### 3.4. Genomic features of two P. multocida strains

Two strains of *P. multocida*, initially identified through ribosomal gene (16S rRNA) sequencing, biotyping, and RAPD grouping with MDR and SBF phenomena were undergone to WGS. The 16S rRNA gene-based phylogenetic analysis revealed that these two *P. multocida* strains had 99.9% identity to Razi_Pm0001 (GenBank accession number: NZ_CP017961.1), and clustered with previously identified *P. multocida* strains in the 16S rRNA phylogeny (Fig. S3). Previous investigations have also shown that *P. multocida* associated with FC represented multiple clones [4, 78]. We found 34 and 37 contigs in PM4 and PM7 strains of *P. multocida*, and the average genome length of these two strains were 2,408,286 bp and 2,408,436 bp, respectively (Table 3). The average GC content of each genome was 40.4% (Table 3), which was consistent with that of a complete *P. multocida* chromosome [8, 16]. The genome completeness of both strains (PM4 and PM7) was 99.55% with genome coverage 458 X and 455.0X for PM4 and PM7, respectively (Table 3). The statistics of the assembly and annotation from NCBI Prokaryotic Genome Annotation Pipeline, Rapid Prokaryotic Genome Annotation (Prokka) [54], and Rapid Annotation using Subsystem Technology (RAST) [55] are summarized in Table 4. According to NCBI annotation pipeline, PM4 and PM7 genomes contained 2260 and 2261 coding sequences (CDS) sequentially where 2217 and 2221 were protein-coding genes (Table 4). Moreover, the number of RNA genes was 58, including 50 transfer RNAs (tRNAs) and 4 rRNAs for each of the genome (Table 4). However, PHASTER database predicted three intact prophages in the genomes of PM4 and PM7 compared with the two intact and one incomplete prophage within the reference strain Razi_Pm0001 (Fig. 4). Noteworthy, despite the difference in biotype and RAPD profiles (Table 2, Fig. S2), both of the strains studied here showed similar genomic features (Table 3). However, PM7 strain harbored two unique genes, and of them, one encoding tonB-dependent hemoglobin/transferrin/lactoferrin family outer membrane receptor facilitating the use of transferrin, lactoferrin, and hemoglobin as sources of iron in different niches within the host [79]. Another unique protein of PM7 was a hypothetical protein of unknown function. Strikingly, endoU domain-containing protein, and inositol-1-monophosphatase genes, involved in inositol metabolism found in both PM4 and PM7 originated *P. multocia* strain, with 100 % aa sequence identity. However, the diversity of inositol fermentation by PM4 (inositol −) and PM7 (inositol +) possibly related to expression of the gene, likely the genes were not expressed properly in PM4, which needs further investigations. These findings indicate all biotypic and RAPD clustering of *P. multocida* does not necessarily directly related to genotypic clusters rather expression of genes plays a determining role in phenotypic classification [80, 81].

**Table 3.**
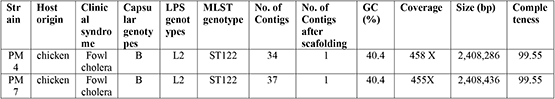
Summary of genome assembly and genotypic characteristics of whole genome sequenced PM4 and PM7 strains of *P. multocida*.

**Table 4.**
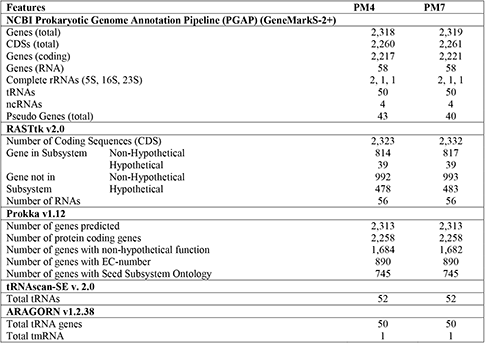
Assembly and annotation statistics of the PM4 and PM7 strains of *P. multocida*.

**Fig. 4.**
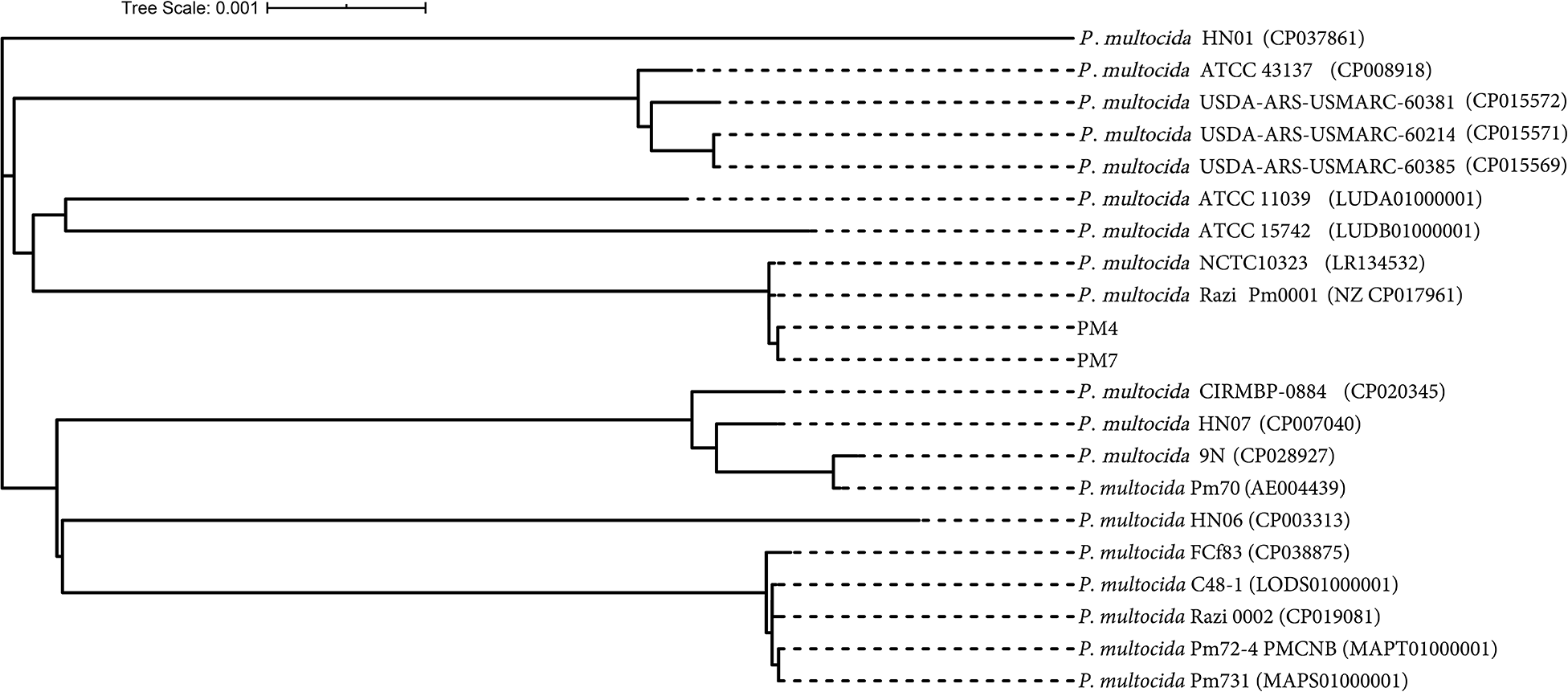
Distribution of virulence factors-associated genes (VFGs), antibiotic resistance genes (ARGs), and prophage related sequence features. Genomes of *P. multocida* strains Razi_Pm0001 and NCTC10323 found to be more closely related to *P. multocida* PM4 and PM7 strains. Capsular serotype determining gene (black), LPS genotyping genes (teal), VFGs (red), ARGs (purple), and prophage (orange) are presented with their respective genomic positions.

### 3.5. Genotypic and phylotypic analysis implicates novel P. multocida type B:L2:ST122 associated with FC

Characterization of the complete genome sequences is an important step in further investigation of the genomic structure, and comparative pathogenomic characteristics of the bacteria. The genome-based phylogenetic tree revealed that *P. multocida* PM4 and PM7 strains clustered in the same branch with the serotype B strains Razi_Pm0001 and NCTC10323 of NCBI of the reference genomes (Fig. 5). The presence of *bcb*D gene in the genomes of PM4 and PM7 determines the capsular serotype B for both isolates PM4 and PM7. The genes *bcb*D is associated with capsular biosynthesis of *P. multocida* specific to serogroup B [7, 11]. On the other hand, rest of the *P. multocida* isolates of the present study showed a higher degree of similarity in RAPD pattern with that of either PM4 or PM7 (Fig. S2) suggesting that these isolates likely to be of the same genotype. These findings are in line with Hotchkiss et al. [69] who reported that closely related *P. multocida* isolates show similar RAPD pattern Moreover, lipopolysaccharide (LPS) outer core structural genes *gat*D, *nct*A, *hpt*F and *hpt*E found in both the isolates indicating LPS genotype L2 as also reported in several earlier studies [7, 17]. Furthermore, based on seven housekeeping genes (*adk*, *est*, *pmi*, *zwf*, *mdh*, *gdh*, and *pgi*) of *P. multocida* in the Bacterial Isolate Genome Sequence Database (BIGSdb), RIRDC MLST type assigned both the PM4 and PM7 isolates into ST122 which is widely documented to be associated with bovine hemorrhagic septicemia (Table 3, Fig. 4). Notably, *P. multocida* from genotype B:L2:ST122 is predominant in bovine [4], and no previous reports show the association of this genotype FC in commercial poultry like laying birds. Moreover, the majority of the avian pasteurellosis outbreaks are caused by *P. multocida* from types A and D [4, 17]. However, cross-species transmission of diseases is frequently reported by different serotypes of *P. multocida* i.e. serogroups A and D. These serotypes are globally distributed, and found to cause diseases in a wide range of domestic animals (e.g. from fowl to calves, pigs, sheep, goats, and rabbits) [15, 17, 22]. Conversely, *P. multocida* serogroups B and E have been found predominantly in tropic areas where they induce hemorrhagic septicemia in cattle and wild ruminants [15, 17, 22]. Moreover, phylogenetically close relatedness of the two isolates of our present study with that of the previously reported bovine originated *P. multocida* type B:L2:ST122. Therefore it is assumed that association of type B:L2:ST122 *P. multocida* with FC might have occurred through host adaptation, and spill-over transmission of the strains from bovine into chicken. Mixed cultivation system of cattle, goat, sheep, and poultry together in the rural areas of Bangladesh as well as poor hygienic practices in the farms further strengthen the possibility of cross-species transmission directly or via the working personnel of the farms [82]. In support with the previous reports, our results indicate that relationship between capsular type, genotype, biotype, host origin, and disease manifestation of *P. multocida* is not clear cut, rather strains from the same capsule, LPS and MLST types can cause similar or different diseases with varied symptoms in multiple hosts despite their geographic origins or phylogenetic relationships [4, 16, 80]. However, further comprehensive studies focusing multi-host adaptation evolution are needed to understand the pathogenesis, mode of transmission, and host adaptability of different serotypes of *P. multocida*.

**Fig. 5.**
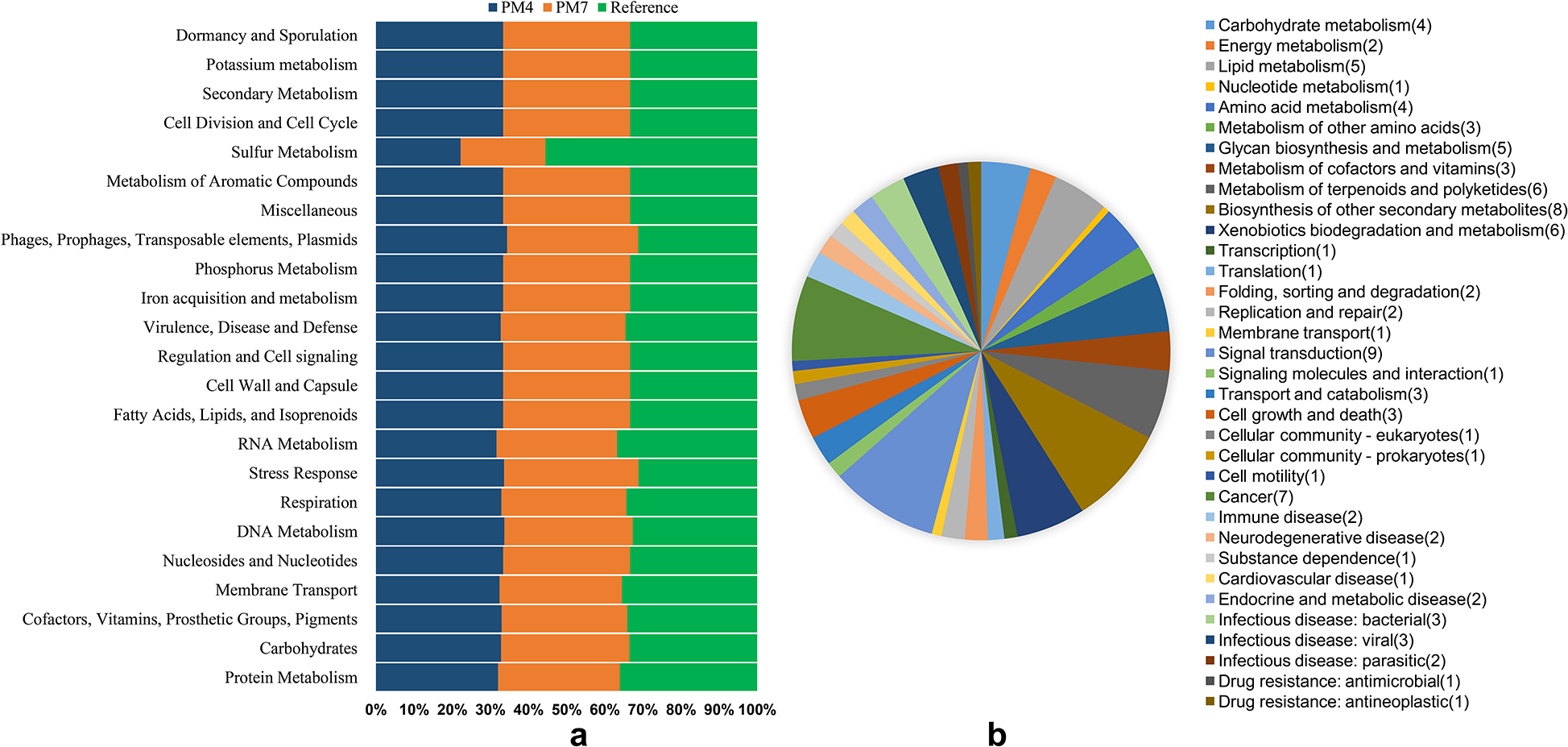
Complete genome-based phylogenetic analysis of *P. multocida* strains. The phylogenetic tree was constructed using the Reference Sequence Alignment Based Phylogeny Builder (REALPHY).

### 3.6. Genome-wide virulence factors-associated genes (VFGs) supporting the pathogenesis of the FC

The genomic features such as coverage, identity and product of different VFGs identified in the complete genomes of the two study strains are in Table 5. In this study, we found different VFGs encoding for outer membrane and porin proteins (*oma*87, *omp*H, *plp*B, *psl*), adhesins (*ptf*A, *fim*A), neuraminidases (*nan*H), iron acquisition related factors (*exb*B, *exb*D, *ton*B, *fur*, *tbp*A, *hgb*A), and superoxide dismutases (*sod*A, *sod*C) in the assembled genome of both PM4 and PM7 isolates (Fig. 4). This result is in agreement with the PCR-based virulence profile of the *P. multocida* strains studied (Table 1, Fig. 4). Potential VFGs were identified based on a homology search against the Virulence Factor Database (VFDB), which identified 14 genes (Table 5) in the PM4 and PM7 genomes, and their pathogenic functions including UDP-3-O-[3-hydroxymyristoyl] N-acetylglucosamine deacetylase, ADP-L-glycero-D-manno-heptose-6-epimerase, UDP-glucose 4-epimerase, D-glycero-beta-D-manno-heptose 1-phosphate adenylyltransferase with 100% query coverage (Table 5). These gene-products are involved in many biosynthetic pathways including secretion system and its effectors, several phospholipases, the elastase and protease IV enzymes, production of phenazines, the exotoxin-A, quorum sensing systems, and synthesis and uptake of the pyochelin siderophore that might have an important role in survival and pathogenesis of *P. multocida* strains avian species [4, 17]. Previous studies reported similar pathogenic gene profiles in *P. multocida* associated with diseases among different hosts including chicken, bovine, porcine and rabbit [4, 11, 16]. Therefore, together these findings indicate neither VFGs nor genotypes can be correlated with host specificity for *P. multocida* [4, 16]. The high frequency of VFGs analyzed was also observed in other studies with strains from both avian and animal hosts [4, 11, 16, 17]. The synchronous distribution of certain VFGs, regardless of host species or *P. multocida* serotype, may suggest the selection of factors that present cross-protection as candidates for vaccine development.

**Table 5.**
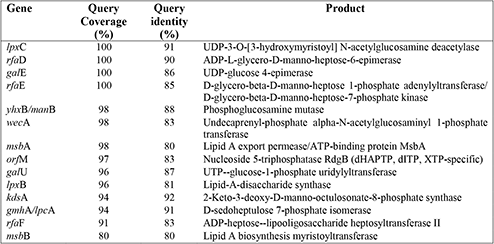
Genome-wide distribution of VFGs, and their features in PM4, PM7 and Razi_Pm0001 strains.

### 3.7. Antimicrobial resistance genes (ARGs) from complete genomes

Comprehensive search for ARGs in the complete genomes of *P. multocida* PM4 and PM7 genomes through the antimicrobial resistance databases such as CARD, ResFinder, and MEGARes explored an array of ARGs (Fig. 4, Table 6). Both PM4 and PM7 genomes seem to be well equipped with a similar load of drug resistance genes conferring resistance to aminoglycoside [*aph*(3’)-Ia, *aph*(3’’)-Ib, *aph*(6)-Id], sulfonamides (*sul*2), tetracyclines (*tet*A, *tet*D), phenicol (*cat*II), trimethroprim (*dfr*), β-lactam (*TEM*-1). The ARGs were categorized into efflux pump conferring antibiotic resistance, antibiotic inactivation enzyme, and antibiotic target in susceptible species (Table 6). The presence of ARGs in the whole genome of PM4 and PM7 corroborated with the resistance pattern observed in antibiotic susceptibility test (Fig. 1, Table 6). In parallel with most Gram-negative bacteria such as *Pseudomonas*, *Vibrio*, *Stenotrophomonas* and *Serratia*, tetracycline resistant *P. multocida* frequently possesses genes *tet*D and *tet*A implicating resistance to tetracycline through efflux of tetracycline or for a protein that prevent tetracycline binding to the bacterial ribosome [25, 83]. Both the strains were resistant to most of the antibiotics tested such as tetracycline, ampicillin, nalidixic acid, oxacillin, cefoxitin and trimethoprim according to EUCAST breakpoints. Although, both the strains, PM4 and PM7 harbored dihydropteroate synthase type-2 (*sul*2), however, the PM4 strain showed complete resistance to the antibiotic sulfonamide, while PM7 exhibited an inhibition zone of 27 mm (Fig. 1). In contrast to our results, Jabeen et al. [25] *pmr*E conferring resistance to polymyxins such as colistin in avian *P. multocida*. Conversely, all of our isolates including PM4 and PM7 were sensitive to colistin corroborating with the absence of *pmr*E gene in their genomes. Remarkably, we did not find any plasmid and integrin representing sequences in the complete genomes of PM4 or PM7 indicating the chromosomal origin of the ARGs [4]. Overall, the abundance of ARGs might be correlated with

**Table 6.**
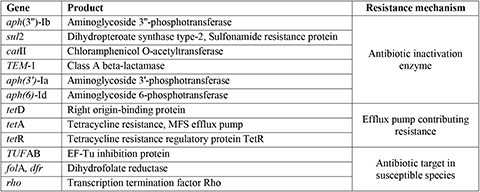
Antimicrobial resistance genes (ARGs) in the genomes of PM4 and PM7 strain.

### 3.8. Metabolic pathways reconstruction

Kyoto encyclopedia of genes and genomes (KEGG) pathway analysis classified the genes of PM4, PM7 and reference strain (Razi_Pm001) into 34 categories, and 356 subsystems (Fig. 6). In these genome, 37% genes were located in the generated subsystems, and the rest were out of the subsystem list. While RASTtk [84] annotation built metabolic pathways of PM4, PM7 and the reference strain related to cellular process, metabolism, environmental information processing, genetic information processing, and pathogenesis (Fig. 6a). Of these subsystems, “amino acids and derivatives” was the largest functional pathway accounting for 190 genes in PM4 and PM7, while the reference strain harbored 199 genes. Similarly, metabolic functional pathways related to “protein metabolism” revealed 175 genes in PM4 and PM7 genomes, and 197in the reference strain. The genome of PM4 harbored 134 genes coding for metabolism of “carbohydrates” whereas 142 and 141 genes were respectively found in the genomes of PM7 and reference strain. In addition, 116 genes associated with “cofactors, vitamins, prosthetic groups, pigments” metabolism were identified in the study genomes (PM4 and PM7) while the reference strain had 120 genes to be related to this metabolic functional pathway (Fig. 6a). The isolates possess several genes associated with adaptation supporting pathways that might help the bacteria being accommodated to pathogenesis in multiple hosts. These pathways include flagellar biosynthesis, motility, quorum sensing, biofilm formation, biosynthesis of vitamin, co-factors, folate, xenobiotics metabolism and so on (Fig. 6b). Diverse metabolic pathways of the bacterial strains reflect their pathogenic fitness, and robust to cause multiple diseases in different hosts [76].

**Fig. 6.**
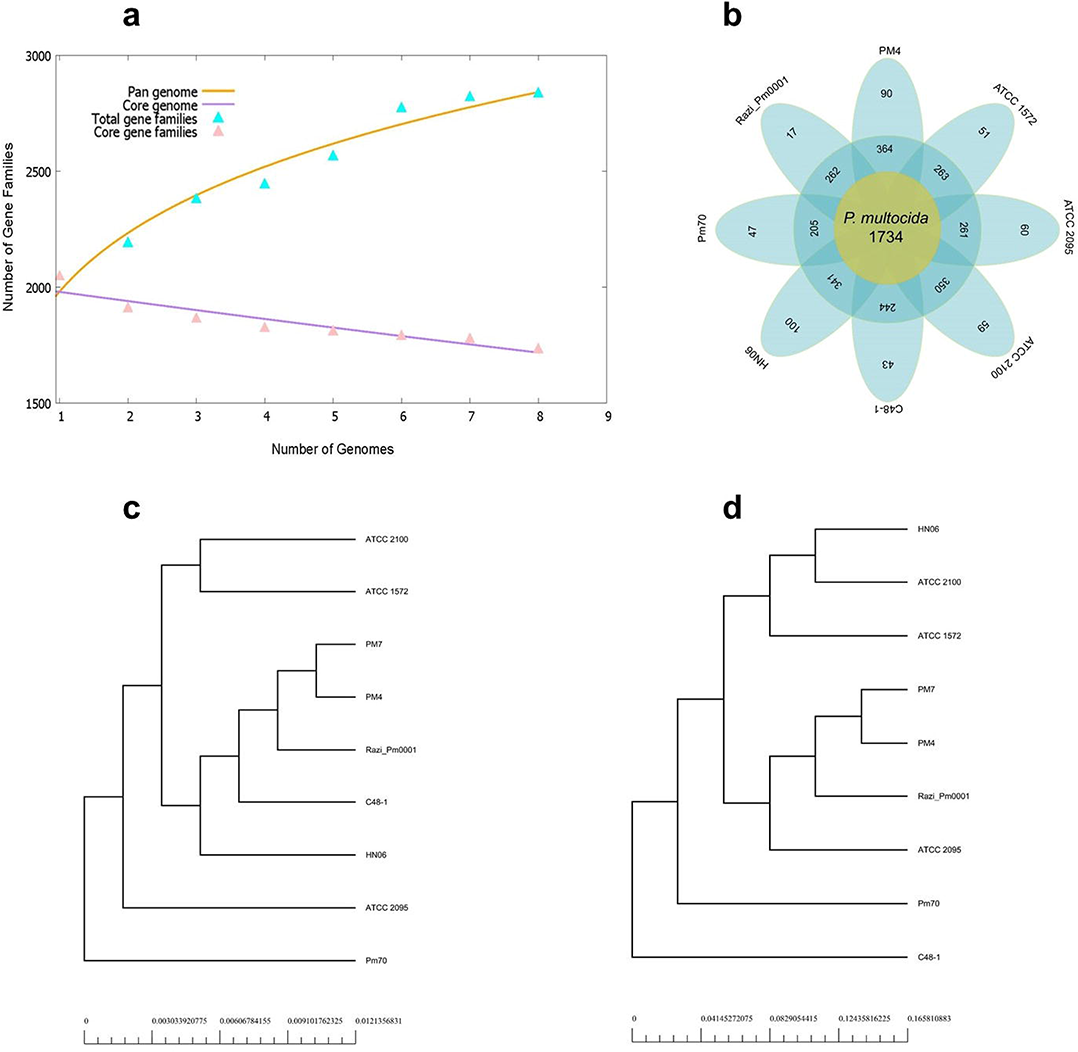
Metabolic pathway reconstruction and subsystem distribution. (a) Comparative gene distribution to different metabolic pathways of PM4, PM7, and the closely related Razi_Pm0001 genomes as predicted by Rapid Annotation System Technology (RAST) Server. (b) The genes classified into 34 categories, and 356 subsystems. The pie chart and count of the subsystem features in the right panel demonstrate the percentage distribution and category of the subsystems found in all the three strains PM4, PM7, and Razi_Pm0001, predicted using KEGG pathway analysis.

### 3.9. Pan-genome analysis of P. multocida

Genome diversity, genome dynamics, species evolution, pathogenesis, and other features of microorganisms have been evaluated by applying pan-genome analysis [85]. The pan genome includes genes present in all strains (core genome), and genes present only in two or more strains of a species (variable or accessory genome). Genes present only in a single strain are called unique genes. To better understand the phylogenetic relationship and bacterial evolution, we performed a pan-genome analysis of seven publicly available whole-genome sequences of *P. multocida* strains with our isolates (Fig. 7, Table S1). The evolution of the pan and core genome is presented in Fig. 7a. In addition, each new genome sequence of *P. multocida*, the number of gene families in the pan-genome increased from 2,131 to 2,838, and that of gene families in the core genome decreased from 1,876 to 1,734 (Fig. 7a, b). The core genome accounted for an average 61.1% of the pan genome. The gene families of the pan genome represent the housing capacity of the genetic determinants and those of the core genome are usually related to bacterial replication, translation, and maintenance of cellular homeostasis [85]. In our study, the unique genes of each *P. multocida* strain exhibited a wide distribution, ranging from 17 (0.6%) to 100 (3.5%). We found 90 unique genes in our isolates (Table S2). These unique genes are found under relaxed mutation pressure, and might have an association with the pathogenicity, virulence and antimicrobial resistance [86], which might have facilitated the strains PM4 and PM7, though being type B:2, to cause FC in layer birds. These types of genes enable the bacteria to transfer benefits to themselves through the horizontal gene transfer, thereby enhancing symbiosis and adaptation of the bacteria to the host, and subsequent onset of pathogenic episodes [87]. However, these potential unique virulence genes need further investigation to prove their association with pathogenicity in FC. The phylogeny based on the pan genome demonstrated that PM4 and PM7 are closer to strain Razi_Pm001 forming a clade with other strains C48-1, and HN06 (Fig. 7c). In contrast, the tree based on the core genome showed that additional to Razi_Pm001 the strain ATCC 2095 belonged to the same clade with our isolates PM4 and PM7 (Fig. 7d). These findings suggest the diverse genetic evolution of the pan and core genomes in different *P. multocida* strains [85].

**Fig. 7.**
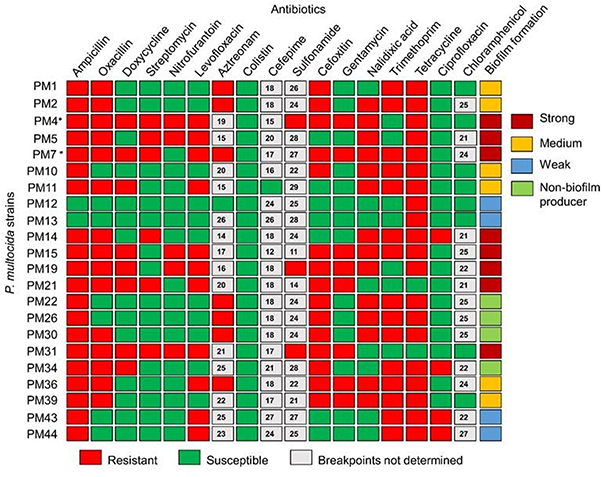
Pan-genome analysis of seven *P. multocida* strains in the repertoire of GenBank. (a) Pan-genome and core genome plot shows the progression of the pan (orange line) and core (purple line) genomes as more genomes are added for analysis. The parameter ‘b’ = 0.17 indicates the pan genome is still open but may be closed soon. The pan genome is still open, as the new additional genome significantly increases the total repertoire of genes. Extrapolation of the curve indicates that the gene families in pan genome increased from 2,131 to 2,838, and those in core genome decreased from 1,876 to 1,734. (b) Flower plot shows the numbers of core genes (inner circle), accessory genes (middle circle), and unique genes (outer circle). (c) Phylogenetic tree based on the pan genome. (d) Phylogenetic tree based on the core genome.

## Conclusions

This study reports the characterization of 22 *P. multocida* isolates from the FC outbreaks in commercial layer farms of Bangladesh. Biochemical and molecular typing grouped the isolates into two major biotypes, and RAPD profiles. The *in vitro* investigation showed that majority of the *P. multocida* isolates were multi-drug resistant, and strong biofilm formers. However, all of the study isolates were sensitive to colistin, suggesting this antibiotic to be effective for treatment of infections in poultry with *P. multocida*. The complete genome analyses of two selected isolates (PM4 and PM7) revealed that the isolates belonged to *P. multocida* genotype B:L2:ST122, which type is commonly found in bovine infections, and not ever reported to cause FC in laying chicken. Furthermore, the comprehensive analysis of the complete genome sequence of *P. multocida* strain PM4 and PM7 has provided valuable insights on the genomic structure with detection and identification of key important genes namely VFGs, ARGs, and genes related to diverse metabolic functions. The pan-genome analysis revealed 90 unique genes involved in versatile metabolic pathways, pathogenicity, virulence and antimicrobial resistance implicating survival fitness and host adaptation advantages among the isolates causing FC. However, further comprehensive genetic and phenotypic study covering multiple hosts required to reveal the genes associated with disease-specificity and host-adaptability of *P. multocida*. Furthermore, a clear phylogenomic relatedness for *P. multocida* can be established based on the complete genome analysis. The insightful findings of this study has opened up an avenue for further research on elucidating the mechanisms behind the molecular pathogenesis of *P. multocida* associated FC in avian hosts with subsequent development of effective preventive and therapeutic strategies.

## Data availability

Complete genome sequences of PM4 and PM7 are available in the NCBI GenBank database under the accession numbers CP052764 (BioSample: SAMN14639261) and CP052765 (BioSample: SAMN14639262), respectively in the BioProject: PRJNA626386.

## Acknowledgments

This work was supported by the grant from Bangladesh Academy of Science – United States Department of Agriculture (BAS – USDA) (Grant no: BAS -USDA PALS DU LSc-34). We would also like to acknowledge Bangabandhu Science & Technology Fellowship Trust for supporting Otun Saha with a PhD fellowship.

## Conflict of interest

The authors declare no conflict of interest regarding this work.

## Ethical approval

The authors confirm the strict adherence to the ethical policies of the journal, as noted on the journal’s author guidelines page. Ethical approval was granted from the Animal Experimentation Ethical Review Committee (AEERC), Faculty of Biological Sciences, University of Dhaka under the Reference No. 71/Biol.Scs./2018-2019.

## Author’s contributions

O.S. carried out the studies (sampling, laboratory experiments, molecular and data analysis). O.S, M.S.R and M.R.I. performed the analyses, and drafted the initial manuscript. M. N. H critically reviewed and interpreted the results, and edited the entire manuscript. M.A.H. and M.S. developed the hypothesis, supervised the work, and critically review the final manuscript. Finally, all authors read and approved the final manuscript.

## Notes

### Competing Interest Statement

The authors have declared no competing interest.

