## Supplementary figures and images for "Genome-wide diversity and differentiation of two novel multidrug-resistant populations of *Pasteurella multocida* type B:2 from fowl cholera"

### Fig. S1

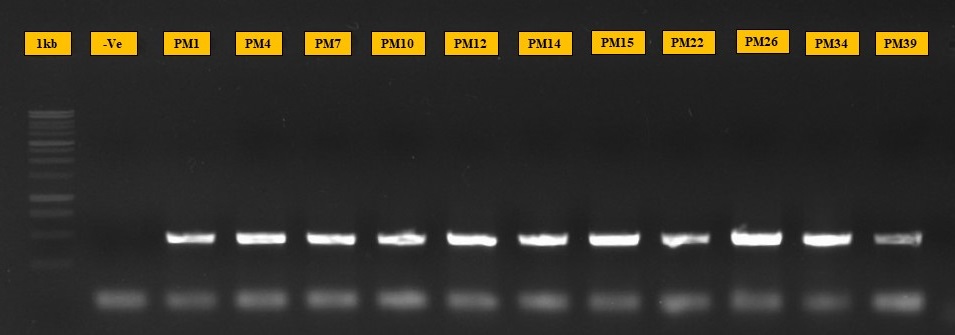

### Fig. S2

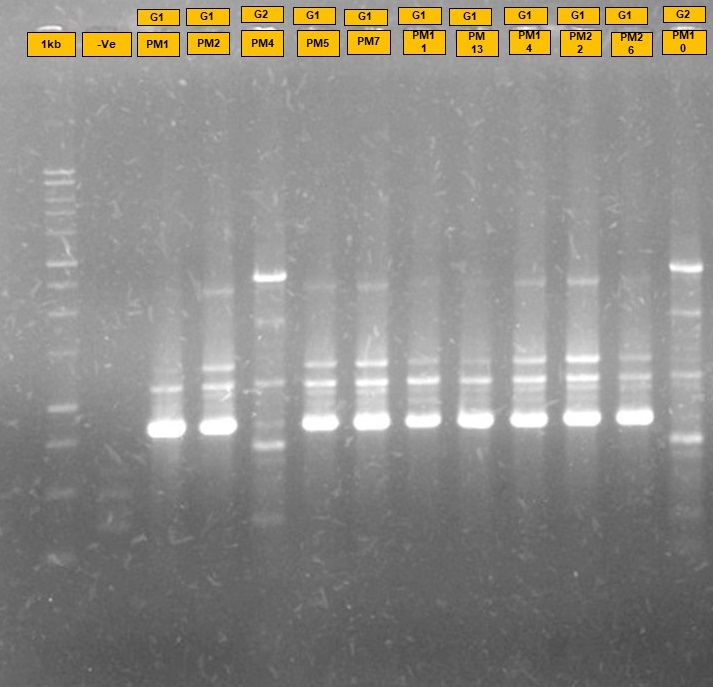

### Fig. S3

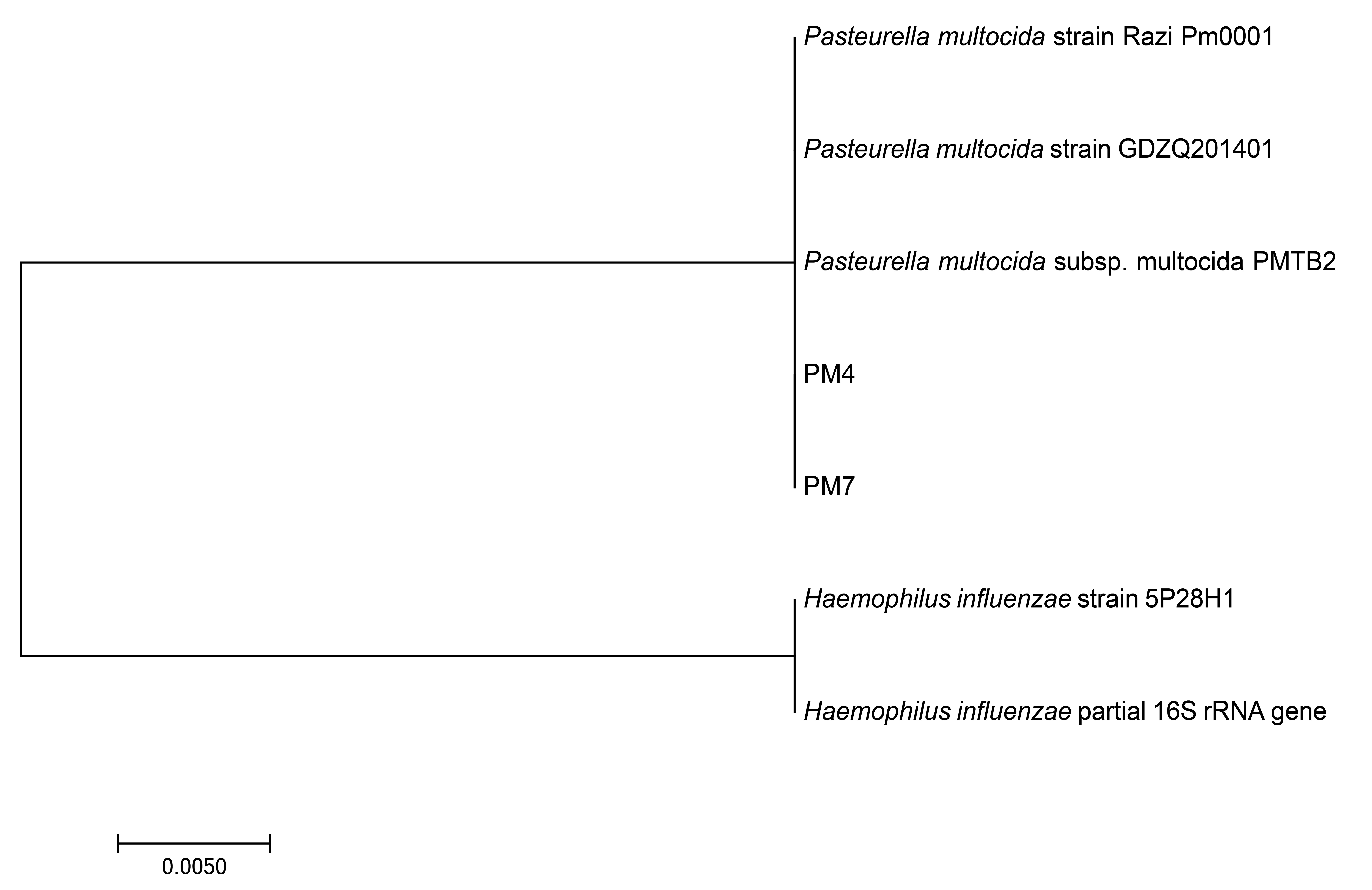
