## Supplementary Tables for "Genome-wide diversity and differentiation of two novel multidrug-resistant populations of *Pasteurella multocida* type B:2 from fowl cholera"

Table S1. Sample information. Twenty-two Pasteurella positive samples used for molecular typing.

| **Isolates** | **Sources** | **Birds** | **Location** |
| --- | --- | --- | --- |
| PM1 | Liver | Live | Narsingdi |
| PM2 | Liver | Dead | Narsingdhi |
| PM4 | Liver | Dead | Narsingdi |
| PM5 | Liver | Live | Narsingdi |
| PM7 | Liver | Live | Narsingdi |
| PM10 | Liver | Dead | Narsingdi |
| PM11 | Liver | Dead | Narsingdi |
| PM12 | Liver | Live | Narsingdi |
| PM13 | Liver | Live | Narsingdi |
| PM14 | Liver | Dead | Narsingdi |
| PM15 | Liver | Live | Narsingdi |
| PM19 | Liver | Dead | Narsingdi |
| PM21 | Liver | Dead | Narsingdi |
| PM22 | Liver | Dead | Narsingdi |
| PM26 | Liver | Live | Narsingdi |
| PM30 | Liver | Live | Narsingdi |
| PM31 | Liver | Live | Narsingdi |
| PM34 | Liver | Dead | Narsingdi |
| PM36 | Liver | Dead | Narsingdi |
| PM39 | Liver | Live | Narsingdi |
| PM43 | Liver | Live | Narsingdi |
| PM44 | Liver | Dead | Narsingdi |
